# Empirically identifying and computationally modelling the brain-behaviour relationship for human scene categorization

**DOI:** 10.1101/2023.01.22.525084

**Authors:** Agnessa Karapetian, Antoniya Boyanova, Muthukumar Pandaram, Klaus Obermayer, Tim C. Kietzmann, Radoslaw M. Cichy

**Affiliations:** Department of Education and Psychology, Freie Universität Berlin, 14195, Berlin, Germany; Charité – Universitätsmedizin Berlin, Einstein Center for Neurosciences Berlin, 10117, Berlin, Germany; Bernstein Centre for Computational Neuroscience Berlin, 10115, Berlin, Germany; Institute of Software Engineering and Theoretical Computer Science, Technische Universität Berlin, 10587, Berlin, Germany; Berlin School of Mind and Brain, Faculty of Philosophy, Humboldt-Universität zu Berlin, 10117, Berlin, Germany; Institute of Cognitive Science, Universität Osnabrück, 49090, Osnabrück, Germany

## Abstract

Humans effortlessly make quick and accurate perceptual decisions about the nature of their immediate visual environment, such as the category of the scene they face. Previous research has revealed a rich set of cortical representations potentially underlying this feat. However, it remains unknown which of these representations are suitably formatted for decision-making. Here, we approached this question empirically and computationally, using neuroimaging and computational modelling. For the empirical part, we collected electroencephalography (EEG) data and reaction times from human participants during a scene categorization task (natural vs. man-made). We then related neural representations to behaviour using a multivariate extension of signal detection theory. We observed a correlation specifically between ∼100 ms and ∼200 ms after stimulus onset, suggesting that the neural scene representations in this time period are suitably formatted for decision-making. For the computational part, we evaluated a recurrent convolutional neural network (RCNN) as a model of brain and behaviour. Unifying our previous observations in an image-computable model, the RCNN predicted well the neural representations, the behavioural scene categorization data, as well as the relationship between them. Our results identify and computationally characterize the neural and behavioural correlates of scene categorization in humans.

**Significance statement:** Categorizing scene information is a ubiquitous and crucial task. Here we provide an empirical and computational account of scene categorization. Previous research has identified when scenes are represented in the visual processing hierarchy, but it remains unclear which of these representations are relevant for behaviour. We identified such representations between ∼100 ms and ∼200 ms after stimulus onset. We then showed that scene categorization in humans can be modelled via a recurrent convolutional neural network in a unified manner, i.e., in terms of neural and behavioural correlates, and their relationship. Together this reveals which representations underlie scene categorization behaviour and proposes a computational mechanism that implements such representations.

## Introduction

Humans effortlessly process visual input from their immediate environment to make adaptively relevant perceptual decisions (Henderson and Hollingworth, 1999). A large body of research has revealed a complex neural cascade involved in processing scenes, emerging across different brain regions (Aguirre et al., 1998; Epstein and Kanwisher, 1998; O’Craven and Kanwisher, 2000; Grill-Spector, 2003; Hasson et al., 2003) and at different time points (Harel et al., 2016; Cichy et al., 2017; Kaiser et al., 2020).

However, it remains unclear which representations are suited to guide behaviour, as identification of activity related to a cognitive function does not imply that this activity can be translated into behaviour. Instead, activity may for example be epiphenomenal (de-Wit et al., 2016), or be related to an interim processing stage that contributes to the creation of representations that later guide behaviour but not do so itself. To identify the representations that are suitably formatted to be used for decision making, behaviour and neural representations must be directly linked (Contini et al., 2017; Grootswagers et al., 2018).

Here we approached this challenge for scene perception from an empirical and a modelling perspective. For the empirical part, we obtained behavioural and brain (EEG) responses simultaneously from participants conducting a scene categorization task. This ensured that the brain measurements directly corresponded to the observed behaviour. We first applied time-resolved multivariate pattern analysis (Cichy et al., 2014; Grootswagers et al., 2017) to reveal the time course with which scene representations emerge over time. To identify the subset suitably formatted for decision making, we then related these unveiled scene representations to the reaction times (RTs) obtained in the scene categorization task. We did so by implementing a distance-to-hyperplane approach (Ritchie and Carlson, 2016) (Fig. 1A), a multivariate extension of signal detection theory (Green and Swets, 1966) previously used to link object representations measured with fMRI (Carlson et al., 2014; Grootswagers et al., 2018) and EEG (Ritchie et al., 2015; Contini et al., 2021) to behaviour. Akin to the criterion in univariate space, it estimates a hyperplane in multivariate space that separates brain measurements for stimuli belonging to two different categories. The distance of the measurements for a stimulus to the hyperplane is assumed to determine behaviour: shorter RTs to the stimulus are associated with longer distances to the hyperplane and vice-versa. Thus, in this framework, a negative correlation between RTs and distances to the hyperplane indicates that the investigated neural representations are suitably formatted for decision-making.

**Figure 1.**
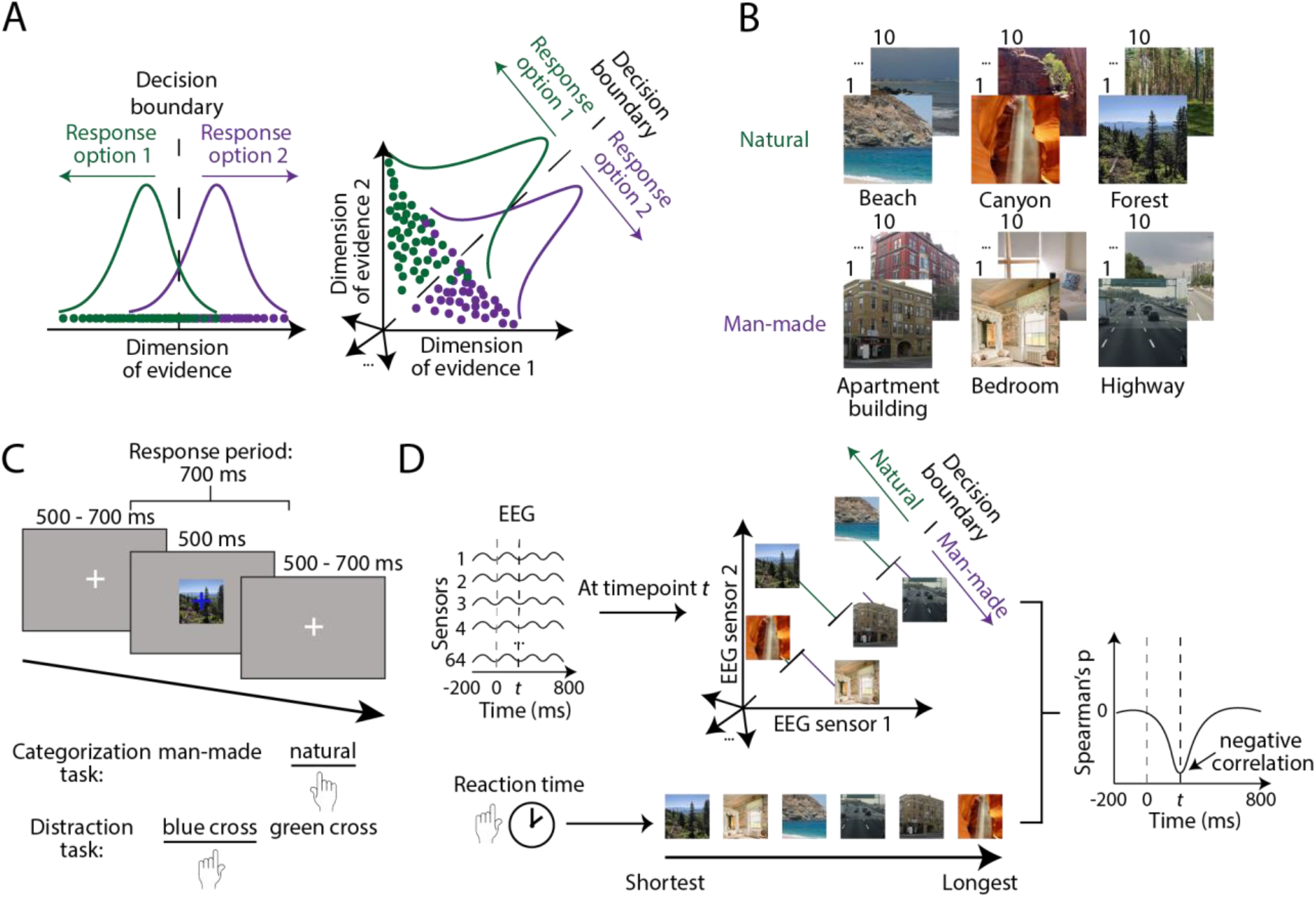
Stimulus set, paradigm and distance-to-hyperplane approach. **A)** Analysis approach. To link neural and behavioural data, we used the distance-to-hyperplane approach, an extension of signal detection theory in multivariate space. **B)** Stimulus set. We selected stimuli from Places-365 (Zhou et al., 2018), creating a set of 30 natural and 30 man-made scenes. **C)** Experimental paradigm. Participants performed a scene categorization task on half of the blocks and an orthogonal fixation cross colour detection task (referred to as distraction task) on the other half. **D)** Distance-to-hyperplane approach. We applied the analysis at every time point to determine when neural representations are suitably formatted for decision-making, which occurs when the correlation between distances and RTs is significantly negative.

Subsequent to linking neural data and behaviour, we used computational models to derive an image-computable model of scene categorization. We consider predictive models an integral part of understanding a scientific phenomenon (see Richards et al., 2019; Lindsay, 2021; Saxe et al., 2021; Doerig et al., 2022b): if we understand a phenomenon, we should be able to provide a model of it that can itself be further evaluated by assessing the importance of model parameters linked to neural parameters (Cichy and Kaiser, 2019).

We formulate three desiderata for a suitable model of scene categorization: it should predict 1) the neural representations underlying scene categorization, 2) human scene categorization behaviour, and 3) their relationship. A potential candidate class for the model are deep convolutional neural networks, which have been shown to predict activity in the visual cortex better than other models (Yamins et al., 2014; Kietzmann et al., 2019; Schrimpf et al., 2020; Cichy et al., 2021). A particular instantiation, a recurrent convolutional neural network (RCNN) named BLnet, i.e., a model with learned bottom-up as well as lateral connectivity, has been shown to predict reaction times in an object categorization task well and better than a range of control models (Spoerer et al., 2020). Based on this observation, we evaluated BLnet with respect to its prediction of human visual scene representations, scene categorization reaction times, and their relationship when relating the behaviour of the model to human representations via the distance-to-hyperplane approach.

## Materials and Methods

### Participants

30 healthy participants took part in the present study (mean age 22.7, *SD* = 2.82; 20 female, 10 male). All participants had normal or corrected-to-normal vision. All participants provided their informed consent after getting acquainted with the study protocol. The study was approved by the ethics committee of Freie Universität Berlin.

### Stimulus set

The stimulus set was composed of 60 scene images selected from the validation set of Places-365 (Zhou et al., 2018). They were center cropped and resized to 480×480. The set contained 30 natural and 30 man-made scenes (Fig. 1B), each of the categories further divided into 3 categories of 10 stimuli (natural: beach, canyon, forest; man-made: apartment building, bedroom, highway).

### Experimental design

Participants were presented with scenes in a random order, overlaid with a green or blue (randomly assigned) fixation cross, on a grey screen (Fig. 1C). On each trial, the scene and fixation cross were presented for 500 ms and were followed by a jittered intertrial interval between 500 ms and 700 ms, where a grey screen and white fixation cross were shown.

Participants performed one of two separate tasks, categorization and fixation cross colour identification (referred to as distraction) (Fig. 1C). They had 700 ms from stimulus onset to report their answer with a button press. In the categorization task, participants had to indicate whether a presented scene was natural or man-made. In the distraction task, participants had to report the colour of the overlaid fixation cross. The key mapping was reversed on every block.

Categorization and distraction trials were presented in alternating blocks (Fig. 1C) of which there were 20 in total. One half of the participants started with one task and the second half started with the other.

For the duration of the experiment, participants were asked to refrain from blinking during the scene trials and to only blink during trials on which a paperclip was shown, which for this purpose was presented for 1000 ms. Paperclip trials were regularly interspersed between main trials every 3-5 trials. Paperclip trials were not included in the analysis.

To ensure that enough data (>20 trials per scene) were available for each participant, every incorrect trial was repeated in the next block of the same task. Therefore, each block contained 3 trials per scene plus the scene trials that were misclassified in the previous block of the same task, as well as paperclip trials which constituted ¼ of all trials in a block.

Right before starting the data collection, participants performed two blocks of the paradigm containing 10 trials each to familiarise themselves with the paradigm.

The experiment was conducted in MATLAB (2019b) using Psychtoolbox (Brainard, 1997).

### EEG recording and preprocessing

Brain activity was recorded using electroencephalography (EEG) with the Easycap 64-electrode system and Brain Vision Recorder. The electrodes were arranged based on the 10-10 system. The participants wore actiCAP elastic caps, connected to 64 active scalp electrodes: 63 EEG electrodes and one reference (Fz). We sampled the activity with a 1000 Hz rate, which was amplified using actiCHamp and filtered online between 0.03 Hz and 100 Hz.

Offline, we preprocessed EEG data using the FieldTrip toolbox (Oostenveld et al., 2010) in MATLAB (2018b and 2021a). First, we segmented raw data into epochs of 1000 ms, using a pre-stimulus baseline window of 200 ms and a post-stimulus window of 800 ms. Then, we performed baseline correction using the 200 ms pre-stimulus window. We applied a low-pass filter of 50 Hz, after which we downsampled the data to 200 Hz, resulting in 200 time points per epoch, each containing the average over 5 ms. To clean the data from artefacts, we used the automatic artefact rejection algorithm from the FieldTrip toolbox (Oostenveld et al., 2010). Additionally, we manually removed noisy channels and trials (mean number of channels removed = 0.5, *SD* = 0.75, mean number of trials removed = 1, *SD* = 1.97). To control for the different levels of noise in the electrodes, we applied multivariate noise normalisation (MVNN; Guggenmos et al., 2018) by multiplying the data by the inverse of the square root of the covariance matrix of electrode activations from the entire epoch. The output of preprocessing was a time course of trial-wise patterns of electrode activations, which we used to perform the analyses described below.

### Scene identity and scene category decoding

We performed scene identity and scene category decoding on subject-level, trial-wise preprocessed EEG data from the categorization and distraction tasks by running classification analyses using a linear support vector machine (SVM; Vapnik, 2013) with the libsvm toolbox (https://www.csie.ntu.edu.tw/~cjlin/libsvm/), in Matlab (2021a). All four analyses contained three main steps, each performed independently on every time point. First, we transformed our data by averaging over individual trials to create “pseudotrials”. Second, we trained the classifier on a subset of data to predict either the scene identity or the scene category of given pseudotrials. Third, we collected the prediction accuracy of the classifier for the left-out data. After running the decoding analyses on all time points and subjects, and averaging over subjects, we obtained a total of four time-courses, depicting scene identity and scene category decoding results for the categorization and distraction tasks.

First, we transformed the trial-wise preprocessed EEG data into pseudotrials by averaging over groups of trials of the same condition in order to boost the signal-to-noise ratio. To ensure that the training of the classifier was not biased, for each pairwise classification we selected the same number of trials per scene in a random fashion, performing this selection and the rest of the analysis 100 times to make use of as much data as possible. For scene category decoding, this was followed by an extra step of splitting the trials from the natural and man-made categories into two groups, such that one half of all trials was used for training and the other half for testing. We then averaged over trials of the same condition (across 4 trials for scene identity and across 20 trials for scene category) to create pseudotrials. For scene category, this step was only performed for the training set: the testing set remained organised by scenes, since we were interested in scene-specific results for further distance-to-hyperplane analyses.

Second, we trained the classifier to predict stimulus conditions using a number of pseudotrials (all but one for scene identity and all training pseudotrials for scene category). We trained the classifier to distinguish between patterns associated with different scenes in scene identity decoding (iterating over all pairwise combinations of scenes) or with the natural/man-made categories in scene category decoding.

Finally, we tested the classifier using the left-out data to assess its prediction accuracy. We presented the classifier with data from two different conditions (depending on the analysis, either different scenes or scenes from different categories), to which it attributed condition labels, and recorded the accuracy of the prediction.

Performing this three-step analysis on all time points and all subjects, and averaging over subjects, resulted in a total of four time-courses of the processing of neural representations associated with scene identity and scene category, in the categorization and distraction tasks.

### Distance-to-hyperplane analysis

The distance-to-hyperplane analysis was performed using the following approach (Fig. 1D). We took the natural/man-made hyperplane estimated during category decoding and calculated, using the same left-out data, the distances to the hyperplane via decision values, which are a unitless measure provided as an output during SVM classification whose absolute value provide information about how close or far scene representations are from the hyperplane.

We then correlated the subject-level distances of all scenes with their RTs (median over subjects) using the Spearman’s rank-order correlation, at every time point, which resulted in one brain-behaviour correlation time course per subject. After averaging over subject-level time courses, we identified the time points when the correlation was significantly negative, which reveal when scene representations are suitably formatted to be used in decision-making.

### EEG channel searchlight analysis

To identify the EEG channels whose signals indicated most the presence of representations associated with scene identity and scene category, as well as the ones that are suitably formatted for decision-making, we combined the decoding and distance-to-hyperplane approaches with a searchlight analysis in channel space. Since we could not perform source reconstruction due to the lack of anatomical scans, we cannot identify where exactly the relevant brain activity is coming from. Instead, we make use of the information provided by the searchlight analysis to identify which channels are involved in the representations of interest, allowing us to approximately infer which regions of the brain contribute to the observed effect.

To implement the searchlight analysis, we performed the decoding (see Section 2.6) and distance-to-hyperplane (see Section 2.7) analyses separately on every channel, by taking into consideration the patterns over the channel and its four nearest neighbours. This resulted in topographic maps indicating for each channel the value of the decoding accuracy, in the decoding analyses, or of the correlation between distances to the hyperplane and RTs, in the distance-to-the hyperplane analysis.

### Fine-tuning and feature extraction of RCNN

We fine-tuned the recurrent convolutional neural network (RCNN) BLnet (Spoerer et al., 2020), initially trained on ecoset (Mehrer et al., 2017) for object classification, on Places-365 (Zhou et al., 2018) in Tensorflow (Abadi et al., 2015) for scene categorization. The training and validation sets for the fine-tuning consisted of samples from 80 scene categories (40 natural and 40 man-made), including the six categories from our stimulus set. The training set contained 30 samples per category, in total 2400 images, while the validation set contained 15 images per category, for a total of 1200 images.

After fine-tuning, we fed the network the 60 scenes that were used in the EEG experiment and collected features from three of its ReLU layers (early (1), mid-level (4) and late (7)) at each time step (8 in total), resulting in feature tensors which were further used in a representational similarity analysis with EEG data to compare network and human scene representations.

### Representational similarity analysis (RSA) between EEG and RCNN

We performed representational similarity analysis (RSA; Kriegeskorte et al., 2008) in two steps: first, we constructed representational dissimilarity matrices (RDMs) for EEG and RCNN features, and afterwards, we correlated these RDMs.

### Construction of RDMs

As a first step, we created representational dissimilarity matrices (RDMs), which are matrices containing dissimilarity values for different conditions, for EEG and RCNN separately.

To create the EEG RDMs, we used subject-level preprocessed EEG data to compute correlation distances (1-Pearson’s coefficient) for each pair of scenes. To ensure that we are using an equal amount of data per condition, we first identified for each subject the minimum number of trials per scene and randomly selected that many trials for every scene. Then, we created pseudotrials by averaging over 5 trials for each scene. For each pair of scenes, we computed the correlation between a pair of pseudotrials. We performed this analysis for all pseudotrials and all pairwise combinations of scenes, 100 times with random assignment of trials to pseudotrials in order to select different subsets of trials every time.

Averaging over all permutations and pseudotrials, we obtained one RDM per subject and time point. Since the RDMs are symmetric matrices, we only used the upper triangular matrix for the analysis, (without the diagonal), which we vectorized in preparation for the next step.

To create the RCNN RDMs, we normalized the extracted features across scenes and calculated the correlation distance (1-Pearson’s coefficient) between the features for two scenes for each pairwise combination of scenes, independently for every RCNN layer and time step. The upper triangular matrix of the RDM for each layer and time step was vectorized.

### Correlation of EEG and RCNN RDMs

To compare the representations of humans and the RCNN, we correlated (Spearman’s correlation) the subject-level EEG RDMs and RCNN RDMs. This correlation was performed independently for each subject, at each EEG time point and for each RCNN layer and time step, resulting in one correlation time course per layer, time step and subject. After averaging over subjects, we obtained a time course of similarity between scene representations of humans and RCNN for each of the three layers and each of the eight time steps. The final time courses depicted the median over the eight time steps.

The analysis was performed three times, for all, natural, and man-made scenes, using the parts of the RDM for the respective scenes. For each of these three time courses, we collected the correlation peak latency, which represents the time point when EEG and RCNN representations are most similar.

### Noise ceiling

To determine the ideal correlation of EEG with the model given the noise levels in our data, we computed the noise ceiling (Nili et al., 2014). To calculate its lower bound, we correlated for each subject at each time point their EEG RDM with the average RDM over the rest of the subjects. This resulted in one correlation time course per subject, and the final lower bound was obtained by averaging over all subjects. The upper bound was calculated in a similar manner, except that the average RDM over subjects also contained the RDM of the subject of interest.

### Collection of RCNN RTs

We extracted RCNN RTs according to the procedure described by Spoerer et al. (2020). In general, according to the principles of threshold-based decision-making (Gold and Shadlen, 2007), humans make a decision (e.g., with a button press) once they accumulate enough evidence for a response option, at which point their reaction time is recorded. To collect RTs from a neural network in a comparable way, we can define a confidence level at which we say that the network has accumulated enough evidence to make a confident decision. Then, we calculate the number of time steps required for the network to reach this confidence level, which represents the network’s reaction time. The confidence level is defined here as a Shannon entropy threshold, which we select based on the fit with human data.

In detail, the procedure to collect RTs from the RCNN was as follows: 1) We defined an entropy threshold between 0.01 and 0.1; 2) We fed the 60 scenes used in the EEG experiment to the network in batches and collected the predictions (natural or man-made) from the readout layer, for each scene and each time step; 3) Using these predictions, we calculated the Shannon entropy for each scene and each time step; 4) We collected the RTs, i.e., for each scene, the first time step (between 1 and 8) that reached the entropy threshold. If the entropy was never reached, the reaction time was defined as 9.

We repeated the steps 1-4 of this procedure for 10 linearly spaced thresholds from 0.01 to 0.1. To select the final entropy threshold and the RTs associated with it, we correlated (Spearman’s correlation) network and human RTs for each of the 10 thresholds. This was performed in a cross-validated manner: for each fold (total 30, representing the number of subjects), we left out the RTs from one subject (different subject at each fold) for a further step and correlated the network RTs with the median RTs over the remaining 29 subjects. This correlation was performed for natural and man-made scenes independently. For each fold, the selected entropy threshold and its associated RTs were the ones for which the network and median human RTs correlated the best and for which the network-human correlation was most similar for the natural and man-made scenes. Here, the entropy threshold was the same for all folds (0.02).

### Statistical analysis

To assess the significance of our results, we performed non-parametric statistical tests.

We first created permutation samples using the sign-permutation test by randomly multiplying each subject’s results either by 1 or -1 10,000 times. The p-value of the original data was obtained by calculating the rank of its test statistic (mean divided by standard deviation) with respect to the distribution of the permutation samples.

Then, we controlled for multiple comparisons by adjusting the p-values of the original data for their inflated false discovery rate (FDR) using the Benjamini-Hochberg procedure (Benjamini and Hochberg, 1995) with ⍺=0.01 for decoding and ⍺=0.05 for the rest of the analyses.

The test was right-tailed for all analyses except distance-to-hyperplane. There, we performed a left-tailed test, as our hypothesis only pertained to negative correlations (i.e., brain and behaviour are correlated when shorter distances to the hyperplane are associated with longer RTs, and vice versa).

All peak latencies and their differences were tested for significance by bootstrapping the subject-level data 1000 times with replacement and calculating the 95% confidence intervals.

## Results

### Behaviour

The mean accuracy across participants for the categorization task was 79.2% (*SD* = 6.75) and 87.6% (SD = 5.89) for the distraction task. Since we only performed analyses on correct trials, this resulted in the inclusion of on average 23.2 (*SD* = 6.0) and 26.0 (*SD* = 1.46) trials per scene respectively for each task.

The mean reaction time for the categorization task was 491.6 ms (*SD* = 35.9 ms) and 446 ms for the distraction task (*SD* = 36 ms).

### Scene identity and category decoding are significant from ∼65 ms after stimulus onset until the end of the trial

We began the neural investigation by revealing the time course with which scene representations emerge and develop over time in the brain using multivariate pattern analysis (also known as decoding) (Kamitani and Tong, 2005; Haynes and Rees, 2006; Carlson et al., 2013; Cichy et al., 2014). This had two aims: we first wanted to ensure that our measurements captured a rich set of candidate representations that could in principle be used by the brain for decision-making, and second, that our results were comparable to previous research, affording theoretical generalisation of our results. For each participant and task context separately, we conducted two classification analyses: scene identity and scene category decoding for the natural vs. man-made division, to reveal, respectively, the time course with which single images and scene categories were distinguishable by their neural representations.

We observed that for both scene identity and scene category decoding (Fig. 2A and Fig. 2B, respectively), regardless of task, the representations of different conditions emerged starting from ∼65 ms after stimulus onset and the conditions remained separable until the end of the trial (p<0.01, FDR-adjusted for each analysis in this section). To visualise how the representations of scenes are distinguished in the brain, we performed multidimensional scaling (MDS) on the results at peak scene identity (Fig. 2C) and category (Fig. 2D) decoding for each task. The plots demonstrate that the representations of natural and man-made scenes at peak decodability are clearly separable in both tasks.

**Figure 2.**
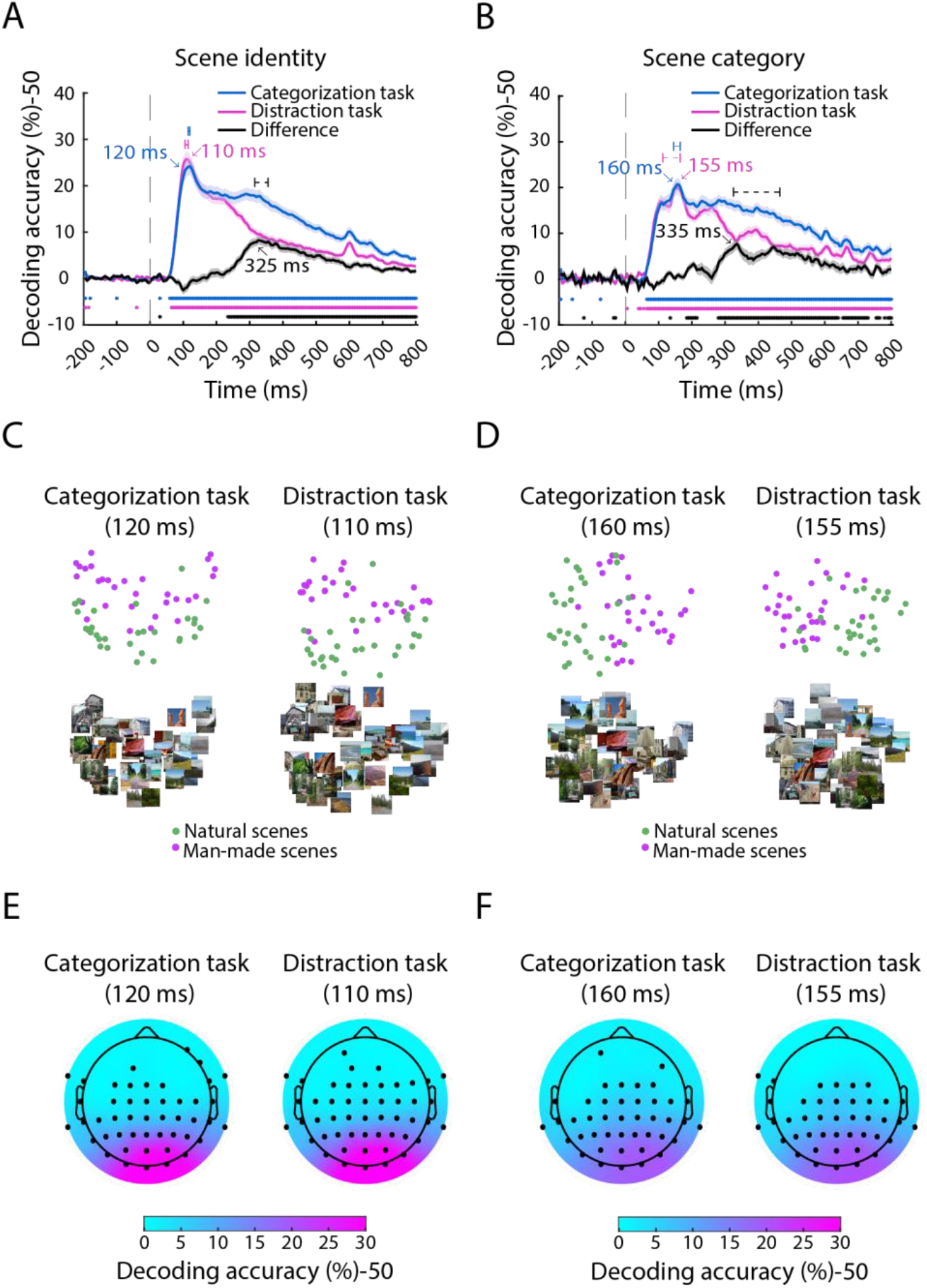
Scene and category decoding. **A)** Pairwise scene identity decoding results on EEG data from the categorization task (blue), distraction task (magenta) and their difference (black). The shaded area around the curves indicates the standard error of the mean (SEM). The dashed horizontal bars represent the bootstrapped 95% confidence intervals of peak latency. The vertical dashed grey line at 0 ms represents the stimulus onset. Significant time points (right-tailed, p<0.01, FDR-adjusted) are indicated with asterisks. **B)** Scene category decoding (natural vs. man-made) results for both tasks and their difference. **C)** MDS results for scene identity decoding from the categorization and distraction tasks at the scene identity decoding peak and **D)** scene category decoding peak. **E)** Results from the searchlight analysis performed in channel space in both tasks at peak decoding latency for scene identity decoding and **F)** scene category decoding. Significant channels (right-tailed, p<0.01, FDR-adjusted) are depicted with black dots.

Further inspection revealed a pattern of results concurrent with previous research in several key aspects. First, scene identity decoding peaked significantly earlier than natural vs man-made category decoding (∼115 ms vs. ∼160 ms, 95% CI of the peak latency difference, averaged over tasks: [15 52.5]), independently of task (for further details, see Table 1) (Carlson et al., 2013; Cichy et al., 2014; Iamshchinina et al., 2022). Second, the time course for categorization and distraction tasks was similar during the early time points and at peak decoding but diverged around 200 ms, independent of the decoding scheme. This suggests that the task impacts visual decoding only after the first feedforward sweep of information processing (Harel et al., 2014; Hebart et al., 2018). Third, a searchlight analysis in channel space (Fig. 2E and Fig. 2F) revealed the topography of the electrodes carrying most information in the scene identity and category classification at the time of peak decoding. We observed a clear focus on occipital electrodes, suggesting that the neural sources of the relevant signals are likely, as expected, in the visual cortex (Cichy et al., 2017; Graumann et al., 2022).

**Table 1.**
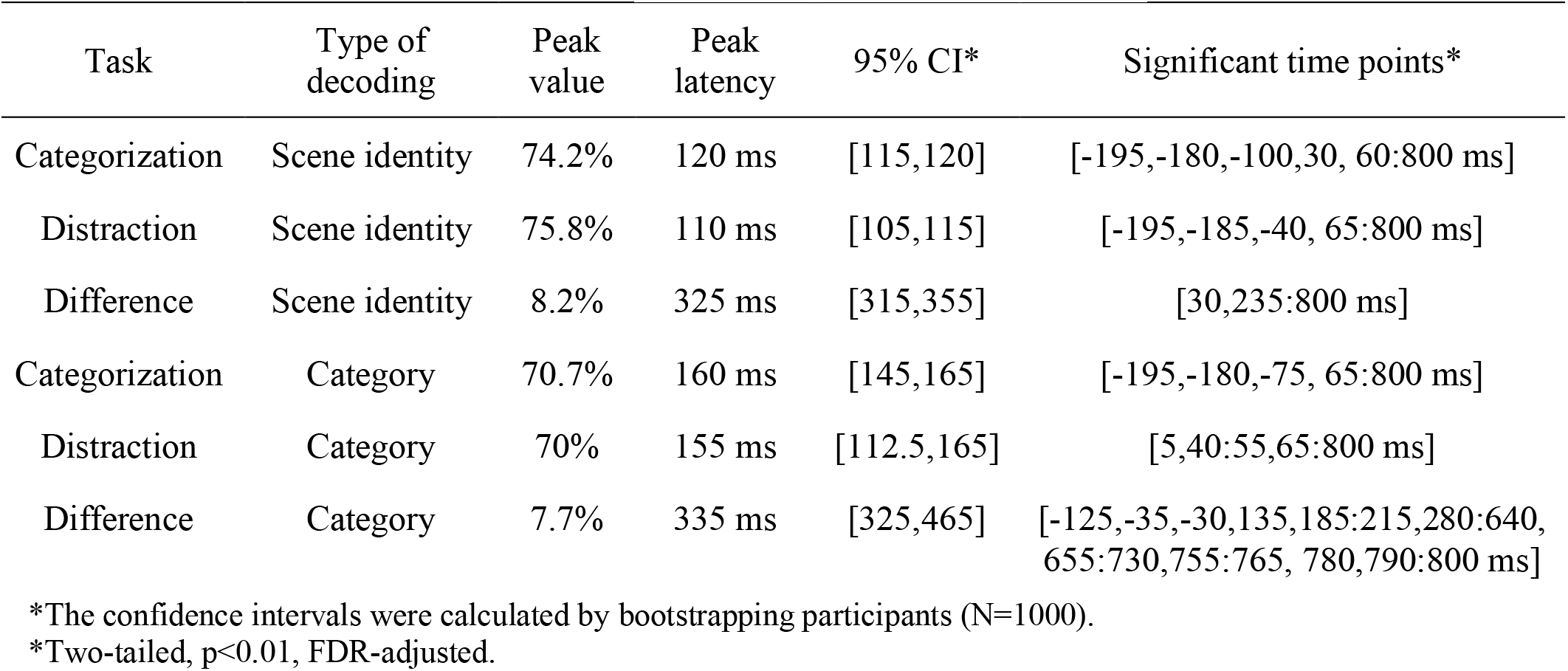
Statistical details for scene identity and scene category decoding.

| Task | Type of decoding | Peak value | Peak latency | 95% CI* | Significant time points* |
| --- | --- | --- | --- | --- | --- |
| Categorization | Scene identity | 74.2% | 120 ms | [115,120] | [-195,-180,-100,30, 60:800 ms] |
| Distraction | Scene identity | 75.8% | 110 ms | [105,115] | [-195,-185,-40, 65:800 ms] |
| Difference | Scene identity | 8.2% | 325 ms | [315,355] | [30,235:800 ms] |
| Categorization | Category | 70.7% | 160 ms | [145,165] | [-195,-180,-75, 65:800 ms] |
| Distraction | Category | 70% | 155 ms | [112.5,165] | [5,40:55,65:800 ms] |
| Difference | Category | 7.7% | 335 ms | [325,465] | [-125,-35,-30,135,185:215,280:640, 655:730,755:765, 780,790:800 ms] |

\*The confidence intervals were calculated by bootstrapping participants (N=1000).
\*Two-tailed, $p < 0.01$ , FDR-adjusted.

Altogether, we verified using decoding that our data yield a temporal results pattern comparable to previous studies, and capture a rich set of candidate representations potentially useful for decision-making. This forms a robust and experimentally well-anchored basis for our further investigation of the link between scene representations and behaviour.

### Distances to the hyperplane in neural space and categorization RTs are negatively correlated between ∼100 ms and ∼200 ms after stimulus onset

To determine when scene information encoded in neural representations is suitably formatted to be used for decision-making, we employed the distance-to-hyperplane approach (Ritchie and Carlson, 2016). This analysis consists of three steps, performed independently at every time point (Fig. 1D): first, we estimated a natural/man-made hyperplane in neural space. Second, we collected the distances of scenes to this hyperplane, and third, we correlated these distances with RTs to the same scenes (but different trials) from a natural/man-made categorization task. Applying the logic of signal detection theory (Green and Swets, 1966) to the neural space, we can identify the points in time at which scene representations and behaviour are statistically linked. The rationale is that distance to a category criterion (here, the categorization hyperplane, estimated in a cross-validated way using a subset of trials) should predict reaction time: stimuli that are harder for humans to categorise should be associated with longer RTs and shorter distances to the hyperplane. Thus, a negative correlation between RTs and distances to the hyperplane in neural space links brain and behaviour.

We performed the distance-to-hyperplane analysis on data from each task separately, as well as across tasks (i.e., using EEG data from categorization and RTs from distraction, and vice versa), to determine the role of the task. We analysed the data here and in subsequent analyses in three subsets: all scenes (Fig. 3A), for a grand-average view; and natural and man-made scenes separately (Fig. 3B and C), for a detailed category-resolved view. All three analyses relating EEG data and categorization behavioural data (Fig. 3A, B and C) converged in showing a significantly negative correlation (i.e., longer RTs were associated with shorter distances to the hyperplane, and vice versa) between ∼100 ms and ∼200 ms after stimulus onset, peaking at ∼160 ms with Spearman’s *⍴* ≈ -0.2 (left-tailed, p<0.05, FDR-adjusted for each analysis in this section; see Table 2 for details). This demonstrates that neural representations arising during this time period are suitably formatted to act as basis for decision-making. Further, it shows that these representations arise independently of whether the relevant categorization task or an unrelated task is carried out, indicating automaticity of the underlying processing (Harel et al., 2014).

**Table 2.** Statistical details from the distance-to-hyperplane analysis.

| Task | Condition | Peak value | Peak latency | 95% CI* | Significant time points* |
| --- | --- | --- | --- | --- | --- |
| Within-task: categorization | All | $\rho=-0.22$ | 160 ms | [160,165] | 140-180 ms |
| Within-task: categorization | Natural | $\rho=-0.21$ | 160 ms | [155,165] | 150-170, 600 ms |
| Within-task: categorization | Man-made | $\rho=-0.25$ | 165 ms | [125,175] | 90-95, 140-180 ms |
| Within-task: distraction | All | - | - | - | - |
| Within-task: distraction | Natural | - | - | - | - |
| Within-task: distraction | Man-made | - | - | - | - |
| Cross-task: categorization EEG, distraction RTs | All | - | - | - | - |
| Cross-task: categorization EEG, distraction RTs | Natural | $\rho=-0.12$ | 130 ms | [-77.5,355] | 125-130 ms |
| Cross-task: categorization EEG, distraction RTs | Man-made | - | - | - | - |
| Cross-task: distraction EEG, categorization RTs | All | $\rho=-0.17$ | 165 ms | [150,175] | 145-175 ms |
| Cross-task: distraction EEG, categorization RTs | Natural | $\rho=-0.16$ | 165 ms | [150,615] | 150-175, 610-615 ms |
| Cross-task: distraction EEG, categorization RTs | Man-made | $\rho=-0.18$ | 105 ms | [100,380] | 75, 90-105, 140-175, 225, 250, 380, 395 ms |

\*The confidence intervals were calculated using bootstrapped permutation samples (N=1000).
\*Left-tailed, $p<0.05$ , FDR-adjusted.

**Figure 3.**
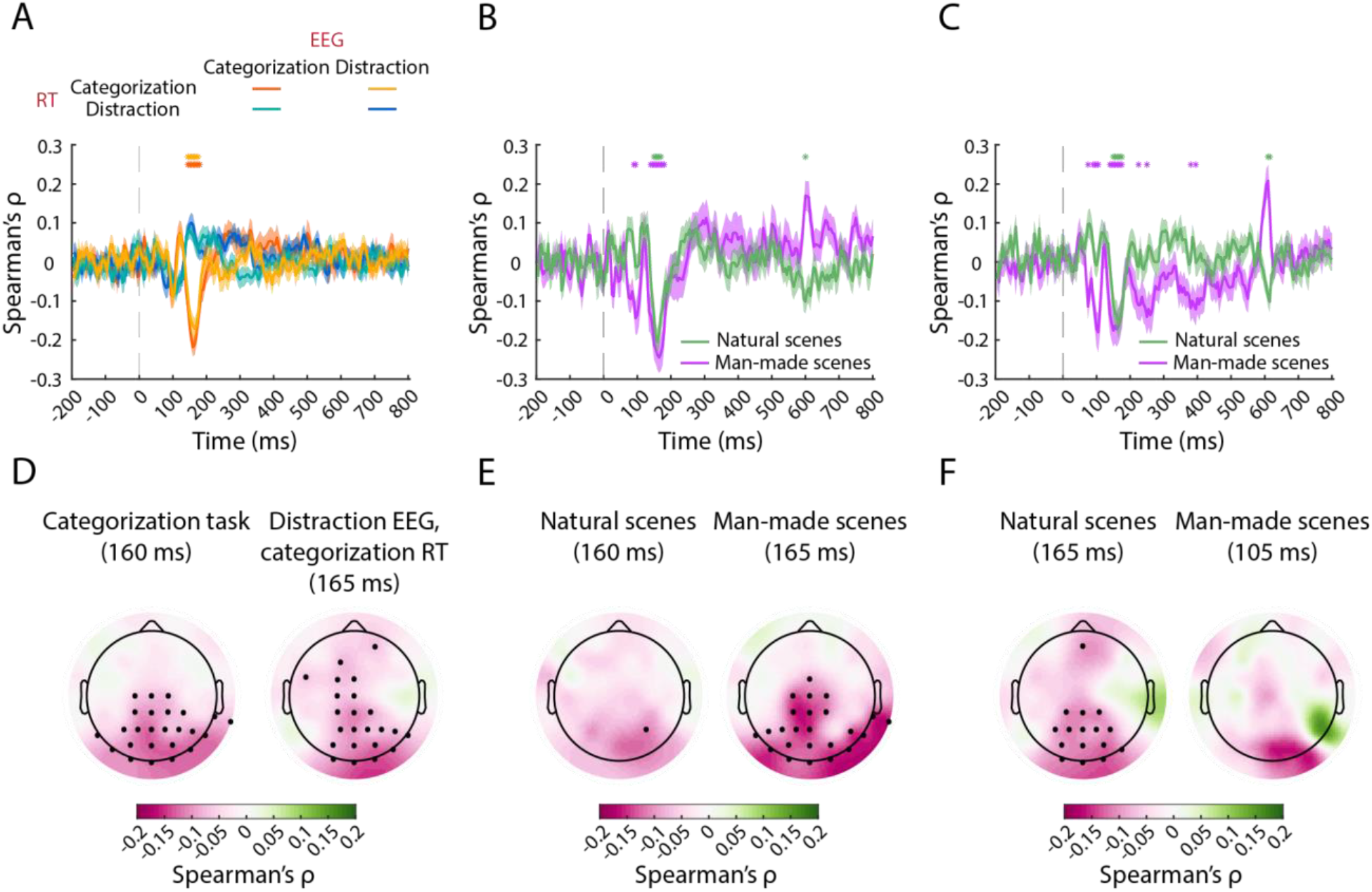
Distance-to-hyperplane analysis. **A)** Results from the analysis on all 60 scenes, on data from the categorization task (orange), the distraction task (blue), using EEG from distraction and RTs from categorization (yellow), and EEG from categorization and RTs from distraction (turquoise). The vertical dashed grey line at 0 ms represents the stimulus onset. The shaded areas around the curves represent the SEM. Significant time points are denoted with asterisks (left-tailed, p<0.05, FDR-adjusted). **B)** Results from the within-categorization analysis on natural (green) and man-made (purple) scenes. **C)** Results from the cross-task analysis using distraction distances and categorization RTs. **D)** Results from the searchlight analysis performed in channel space at peak negative correlation latency on all scenes in the categorization task (left) and cross-task using EEG data from distraction and RTs from categorization (right). **E)** Results from the searchlight analysis on natural and man-made scenes in the categorization task and **F)** the cross-task analysis with EEG from distraction and RTs from categorization. The negative correlations are in pink. Significant channels (left-tailed, p<0.05, FDR-adjusted) are depicted with black dots.

For the control analyses relating EEG data with RTs from the scene category-unrelated distraction task (Fig. 3A, blue and turquoise curves), there were no significant negative correlations between distances and RTs. This ascertains the specificity of the identified link between scene representations and classification behaviour.

To identify the EEG channels whose signals indicated most the presence of representations that are suitably formatted for decision-making, we combined the distance-to-hyperplane approach with a searchlight analysis in channel space. In detail, we conducted one searchlight analysis (Fig. 3D, E, and F) for each distance-to-hyperplane analysis described above (Fig. 3A, B, and C) at the latency of the respectively identified peak. Consistent across all three analyses, we observed the strongest negative correlations in the occipital electrodes, suggesting the origin of the identified behaviourally-relevant representations to be in the visual brain.

Interestingly, we observed that in analyses involving EEG from the distraction task (Fig. 3D, right, and Fig. 3F, left), a significant effect arose also in anterior electrodes overlying the frontal cortex. This might suggest that in contexts where automatic categorization is hindered by an unrelated task, frontal brain regions may contribute to representations of visual category suitably formatted for decision making, consistent with the role of frontal cortex in processing object representations (Freedman et al., 2001, 2003; Bar, 2003; Bar et al., 2006; Kar and DiCarlo, 2021). In sum, our results indicate that scene representations emerging automatically in the visual brain between ∼100 and ∼200 ms after stimulus onset are suitably formatted to be used for decision-making.

### Humans and RCNNs correlate in terms of neural representations, RTs and the relationship between representations and RTs

Based on the empirical work, we aimed at providing a computational model of scene categorization in humans. As a candidate model, we selected BLnet (Spoerer et al., 2020), a recurrent convolutional neural network that has specifically been shown to predict RTs to objects, and that, as a deep neural network trained on an object classification task, belongs to the class of models that predict human visual cortex activity well (Kar et al., 2019; Kietzmann et al., 2019; Schrimpf et al., 2020; Cichy et al., 2021).

The model consists of seven layers and contains bottom-up as well as lateral recurrent connections at every layer (see Fig. 4A for architecture details). These lateral connections provide features from eight different time steps at each layer. The model was trained on object classification, and as training material specificity has been shown to impact predictive power for brain representations (Yamins et al., 2014; Cichy et al., 2016, 2017; Mehrer et al., 2017), we first fine-tuned all its layers on a scene categorization task (natural vs. man-made) using a database of scenes (Places-365, Zhou et al., 2018).

**Figure 4.**
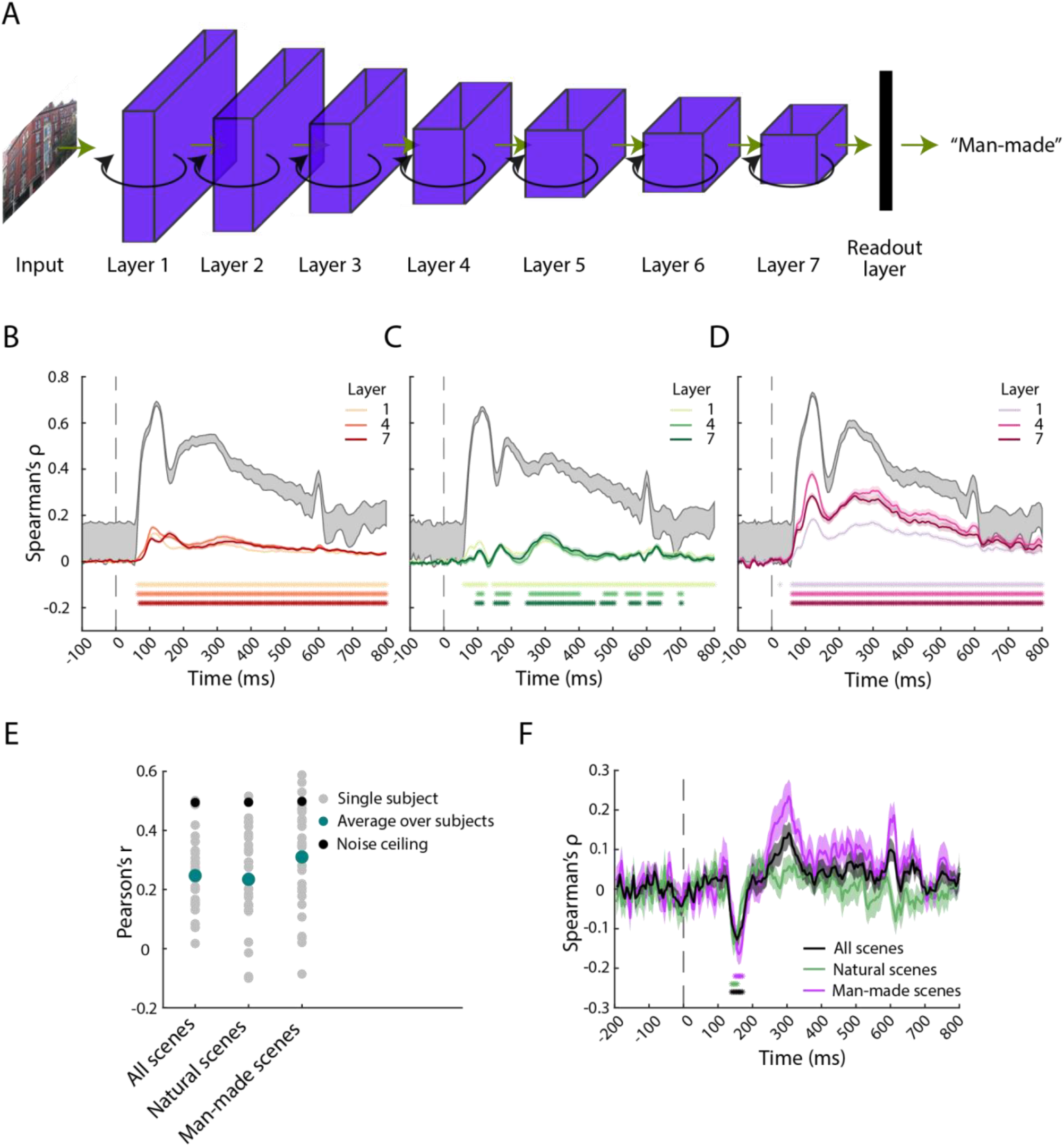
Modelling human visual scene categorization with an RCNN. **A)** Architecture of BLnet (Spoerer et al., 2020), the recurrent DNN used in the analysis. The network consists of seven layers, linked via bottom-up (green arrows) and lateral (black arrows) connections. Features were extracted from three layers (1, 4 and 7) at eight different time steps, and RTs were collected from the readout layer. **B)** Results of the representational similarity analysis performed on the neural representations of humans and the RCNN features from three different layers (median over eight time steps), for all scenes, **C)** natural scenes and **D)** man-made scenes. The vertical dashed grey line at 0 ms represents the stimulus onset. The shaded areas around the curves represent the SEM. Significant time points are denoted with asterisks (right-tailed, p<0.05, FDR-corrected). The shaded grey area represents the noise ceiling. **E)** Results of the correlation between scene categorization RTs of humans and of the RCNN. The correlation is significant for all analyses (right-tailed, p<0.05, FDR-corrected). **F)** Results of the distance-to-hyperplane analysis performed with distances from EEG data and RCNN RTs. Significant time points are denoted with asterisks (left-tailed, p<0.05, FDR-adjusted).

We assessed three desiderata for a model of scene categorization: the model should predict 1) the neural representations underlying scene processing, 2) human behaviour, and 3) their relationship. For the first desideratum, we compared human and BLnet representations using representational similarity analysis (RSA; Kriegeskorte et al., 2008). For the model, we focused on three network layers: early (layer 1), mid-level (layer 4) and late (layer 7) as representatives for respective visual processing stages, and performed the analysis on all eight time steps. For the EEG, we built RDMs in a time-resolved fashion for every time point containing the average of five milliseconds. Comparing RDMs (Spearman’s *⍴*) yielded time courses where positive correlations indicate similar scene representations in humans and BLnet.

Consistent across the analyses on all (Fig. 4B), natural (Fig. 4C) and man-made scenes (Fig. 4D), we observed, starting from ∼60 ms, significant positive correlations for all of the trial duration (right-tailed, p<0.05, FDR-adjusted for each analysis in this section). Assessing the model’s different time steps yielded comparable results, therefore Fig. 4B, C, D depicts the median over all time steps.

Focusing in detail on the analysis of all scenes (Fig. 4A), we observed a forward shift in peak latency with increased network depth. In layers 1 and 4, the correlation peaked at ∼110 ms, while in layer 7, the peak correlation was significantly later at 160 ms (95% CI of difference between layers 1 and 4, and 1 and 7: 47.5 ms [20 65]; see Table 3 for further details), suggesting a temporal correspondence between network layer and processing stage at which different scene representations emerge (Güçlü and Gerven, 2015; Cichy et al., 2016; Eickenberg et al., 2017; Greene and Hansen, 2018; but see Sexton and Love, 2022). Focusing on the finer distinction between natural and man-made scenes (Fig. 4C vs. D), we observed overall weaker effects for natural scenes. Nevertheless, the results show that the model fulfils the first criterion of predicting human scene representations for all types of scenes.

**Table 3.**
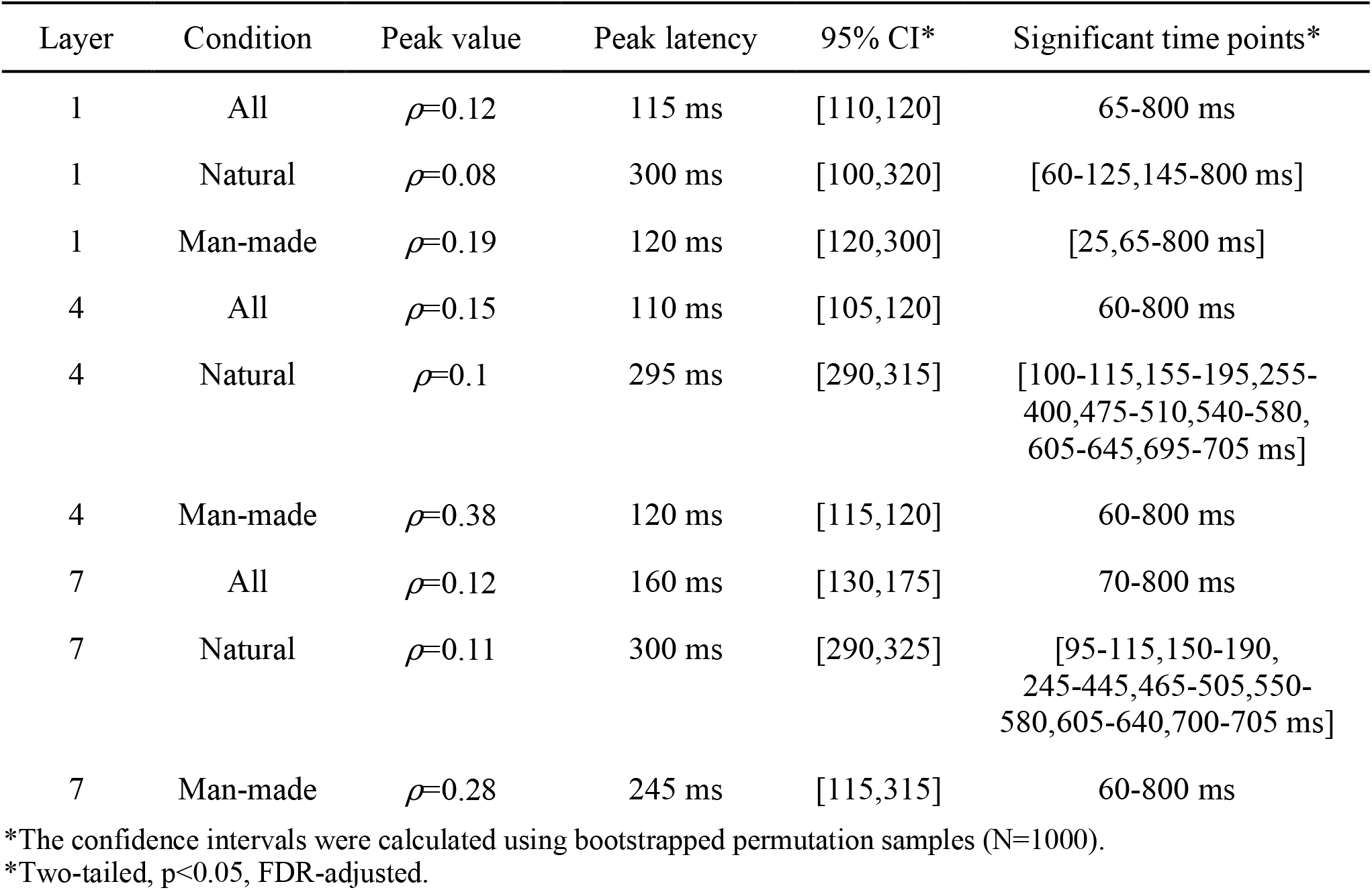
Statistical details for the RSA between humans and the RCNN.

To assess our second desideratum, i.e., the prediction of behaviour in terms of RTs in a categorization task, we compared RTs from BLnet and humans. In short, we extracted BLnet RTs (Spoerer et al., 2020) by fitting an entropy threshold in a cross-validated way and collecting the network time step at which this entropy threshold was reached for each of the scenes, which then served as reaction time. We then correlated these network RTs with human RTs. As before, this analysis was performed separately for all scenes, for natural, as well as for man-made scenes. For all three analyses (Fig. 4E), we observed significant positive correlations (Pearson’s *r* = 0.25, 0.31 and 0.24 respectively; right-tailed, p<0.05, FDR-adjusted), demonstrating that BLnet’s reaction times significantly correlate with human behaviour.

Lastly, we assessed our third desideratum, i.e., whether there is an analogy for BLnet and humans in the relationship between representations and behaviour. For this we conducted the distance-to-hyperplane analysis using human EEG data and BLnet RTs. This analysis is non-trivial, as the correlation between human and network reaction times is not so strong as to automatically suggest a significant correlation between network RTs and EEG.

Nevertheless, we observed that this analysis consistently yielded a significant negative correlation between ∼100 ms and ∼200 ms after stimulus onset (Fig. 4F), analogous to the results from the analysis based on human rather than model RTs (see Fig. 3A, B). This demonstrates that BLnet successfully mirrors the relationship between human visual scene representations and behaviour.

In sum, we find that the model predicts neural representations, reaction times and the link between brain and behaviour, thereby fulfilling all three formulated desiderata for a computational model of visual scene categorization in humans.

## Discussion

We investigated scene processing in humans using multivariate analyses of EEG data, reaction time measurements, and computational modelling based on a recurrent convolutional neural network, BLnet (Spoerer et al., 2020). We highlight two main findings. First, using the distance-to-hyperplane approach on the empirical EEG and behavioural data, we found that neural representations of scenes are negatively correlated with RTs between ∼100 ms and ∼200 ms after stimulus onset, indicating that neural representations are then suitably formatted for decision-making. Second, we demonstrated that RCNNs are good predictors of neural representations, behaviour and the brain-behaviour relationship for scene categorization, fulfilling all three desiderata that we formulated to identify a suitable model of scene categorization in humans.

### Neural representations of scenes are suitably formatted for decision-making between ∼100 ms and ∼200 ms after stimulus onset

Using brain decoding, we revealed that individual scenes and scene categories are represented in the brain continuously starting from ∼65 ms post stimulus, with peaks between 100 ms and 200 ms (Groen et al., 2013; Harel et al., 2016; Cichy et al., 2017; Greene and Hansen, 2020; Kaiser et al., 2020). Resolving the time course of scene representations does not, however, reveal conclusively when these representations are suited for use during decision-making. Performing the distance-to-hyperplane approach, we showed this to be the case in a short time-window between ∼100 ms and ∼200 ms post stimulus, coinciding with peak decoding.

The timing of the brain-behaviour relationship is consistent with previous univariate studies (VanRullen and Thorpe, 2001; Philiastides and Sajda, 2006; Philiastides et al., 2006; Greene and Hansen, 2020) and studies investigating other visual contents using other measurements of behaviour, such as perceived similarity in abstract stimuli (Wardle et al., 2016), objects (Bankson et al., 2018; Cichy et al., 2019), and scenes (Greene and Hansen, 2020). This consistent temporal pattern for different visual contents suggests similar underlying neural mechanisms through which visual representations suitably formatted for behaviour emerge.

We observed the brain-behaviour relationship even for brain data recorded during a visual task unrelated to categorization behaviour (Carlson et al., 2014; Harel et al., 2014; Ritchie et al., 2015; Grootswagers et al., 2018). This is consistent with the view of object categorization as a core cognitive function of the visual system (VanRullen and Thorpe, 2001; Grill-Spector, 2003; DiCarlo et al., 2012). Therefore, this finding highlights the significance and automaticity of categorising perceptual information and of processing it into a behaviourally-guiding format, enabling quick adaptive decision-making.

Interestingly, we observed a relationship between brain measurements and behaviour in frontal electrodes when relating reaction times from object categorization to EEG from the distraction task. While we cannot infer the neural sources of the signal from the results of this analysis, they suggest that representations suited for behaviour could be formed not only in the visual cortex, but also in the frontal regions. This finding would follow (Grootswagers et al., 2018), who observed activity in the prefrontal cortex during an orthogonal task, and is consistent with evidence from previous animal and human studies suggesting that frontal areas relevant for task performance are activated in decision-making (Hanes and Schall, 1996; Kim and Shadlen, 1999; Heekeren et al., 2006, 2006; Philiastides and Sajda, 2007; McGinty and Lupkin, 2021; Stringer et al., 2021). Further research is required to specify the nature and cortical source of this effect.

### RCNN can be used as a suitable and unified model of human scene categorization

We observed a positive correlation between the representations of the network and humans from ∼60 ms post-stimulus until the end of trial, consistent with previous findings in scene (Cichy et al., 2017; Greene and Hansen, 2018) and object (Cichy et al., 2016; Jozwik et al., 2017; Seeliger et al., 2018; Kietzmann et al., 2019) recognition. Our results therefore add evidence towards the idea that RCNNs are good predictors of human neural representations (Cadieu et al., 2014; Khaligh-Razavi and Kriegeskorte, 2014; Yamins et al., 2014; Güçlü and Gerven, 2015), particularly for scenes (Doerig et al., 2022a).

Having similar representations to humans’ is not enough for a network to qualify as a suitable model of human vision: it must also behave similarly to humans. This was the case for BLnet: we observed for the first time a positive correlation between human and network categorization RTs for all types of scenes. Our results contribute to efforts comparing human and DNN behaviour in terms of performance (Seijdel et al., 2020), similarity judgments (Jozwik et al., 2017; King et al., 2019), error consistency (Geirhos et al., 2021) and reaction times (Rafiei and Rahnev, 2022; Sörensen et al., 2022). In particular, we extend previous results relating RCNNs to human behaviour from object recognition (Spoerer et al., 2020) to scene categorization, demonstrating the potential of RCNNs as models for diverse visual human behaviours.

Lastly, we observed that the relationship between EEG distances and network RTs between ∼100 ms and ∼200 ms post stimulus corresponded directly to the empirical results from the within-humans analysis, suggesting that RCNNs can be used to successfully model the brain-behaviour link. It could additionally imply that the representations which are similar in humans and the RCNN are the ones feeding into scene categorization behaviour. This brings the analysis full circle by integrating brain measurements, behaviour and deep networks in a unified modelling account of human perceptual decision making.

However, the modelling results should be interpreted with caution: although the RCNN correlates strongly with humans both in terms of neural representations and behaviour, further research is needed to identify how much variance it explains in human data. There is room for improvement through better models, e.g., by employing different objective functions, adding top-down connections, adding more parameters, and so on, in order to close the gap between computer and human vision (Kietzmann et al., 2019; Geirhos et al., 2021; Zador et al., 2022).

### Limitations and future directions

There are several limitations to our results. First, the distance-to-hyperplane approach operationalizes behaviour only in terms of RTs. It is possible that other neural representations are linked to other aspects of behaviour, e.g., accuracy or similarity judgements at different time points. Further, while we only observed effects between ∼100 ms and ∼200 ms post stimulus, we cannot exclude that the strong signal-to-noise ratio in this time-window influences the result. Using a more precise neural measurement could reveal whether other time-points are also involved in decision-making. Lastly, we only assessed one network and cannot draw any conclusions regarding the usefulness of particular parameters, such as recurrence, architecture, training material, etc. in modelling scene categorization.

Future studies should test other models such as feedforward DNNs to determine the importance of each of those factors.

### Summary

In this work, we explored the link between neural representations and behaviour using empirical and computational methods. Empirically we showed that brain and behaviour are linked during peak brain decoding, i.e., when neural representations of different scenes are most distinguishable. Computationally, we demonstrated that an RCNN is a unified model of scene processing, suggesting that future studies can use RCNNs to further understand scene processing in humans in terms of both neural and behavioural data.

### Code and data availability

The code used for this project can be found under https://github.com/Agnessa14/Perceptual-decision-making.

The data and stimulus set can be found on https://osf.io/4fdky/.

## Acknowledgments

A.K. is supported by a PhD fellowship of the Einstein Center for Neurosciences. R.M.C. is supported by German Research Council (DFG) grants (CI 241/1-1, CI 241/3-1, CI 241/1-7) and the European Research Council (ERC) starting grant (ERC-StG-2018-803370). We thank the HPC Service of ZEDAT, Freie Universität Berlin (Bennett et al., 2020) for computing time, as well as NVIDIA Corporation for the donation of the GPU used for this research. We gratefully acknowledge the contribution of Alessandro Gifford, Greta Häberle, Johannes Singer and Siying Xie and their valuable comments on the manuscript. We would also like to thank Martin Hebart, Youssef Kashef, Thomas Carlson, J. Brendan Ritchie and Tijl Grootswagers for their input on the analysis.

